# A role for a conserved kinase in the transcriptional control of methionine biosynthesis in *Escherichia coli* experiencing sustained nitrogen starvation

**DOI:** 10.1101/254623

**Authors:** Amy Switzer, Dimitrios Evangelopoulos, Rita Figueira, Luiz Pedro S. de Carvalho, Daniel R. Brown, Sivaramesh Wigneshweraraj

## Abstract

The initial adaptive transcriptional response to nitrogen (N) starvation in *Escherichia coli* involves large-scale alterations to the transcriptome mediated by the transcription activator, NtrC. One of the NtrC-activated genes is *yeaG*, which encodes a conserved bacterial kinase. Although it is known that YeaG is required for optimal survival under sustained N starvation, the molecular basis by which YeaG benefits N starved *E. coli* remains elusive. By combining transcriptomics with targeted metabolomics analyses, we demonstrate that the methionine biosynthesis pathway becomes transcriptionally dysregulated in *ΔyeaG* bacteria experiencing sustained N starvation. This results in the aberrant and energetically costly biosynthesis of methionine and associated metabolites under sustained N starvation with detrimental consequences to cell viability. It appears the activity of the master transcriptional repressor of methionine biosynthesis genes, MetJ, is compromised in *ΔyeaG* bacteria under sustained N starvation, resulting in transcriptional derepression of MetJ-regulated genes. The results suggest that YeaG is a novel regulatory factor and functions as a molecular brake in the transcriptional control of both the NtrC-regulon and methionine biosynthesis genes in *E. coli* experiencing sustained N starvation.

---

Conditions that sustain constant bacterial growth are seldom found in nature. Environments where growth is limited by availability of nutrients are common, for example, soil, water, or even host environments such as macrophages can lack essential nutrients to support growth. As such, many bacteria spend most of their time in states of little or no growth due to starvation. The starved and growth attenuated state is now widely considered as an important physiological condition in bacterial pathogenesis and survival. Nitrogen (N) is an essential element of most macromolecules (proteins, nucleic acids and cell wall components) within a bacterial cell. Many bacterial pathogens appear to experience N limitation in host environments such as the urinary tract (e.g. uropathogenic *Escherichia coli* (1)) or macrophages (e.g. *Salmonella* Typhimurium (2)), and respond by activating specific adaptive processes. Emerging evidence also indicates that bacterial N metabolism and N stress responses have a role in the development of dysbiosis (3). Further, we recently reported that the specific adaptive response to N starvation in *E. coli* is directly coupled to the production of guanosine pentaphosphate, (p)ppGpp: the pleiotropic stress signaling nucleotide and effector of the stringent response (4,5). Thus, the adaptive response to N starvation clearly plays an integral and fundamental role in bacterial physiology and pathogenesis.

The initial adaptive response to N starvation is mediated by the transcriptional activator, NtrC, and involves the expression of genes largely associated with activation of transport systems, catabolic and biosynthetic operons for the scavenging for alternative N sources, and managing cellular resources until growth conditions improve. One of the highly expressed operons during the initial adaptive response to N starvation consists of *yeaG* and *yeaH.* Both genes are conserved across several bacterial species, especially Enterobacteriaceae (5-7). They are also induced in response to diverse stresses (including low pH (8), high osmolarity (9), stationary phase (8) and low sulphur (10)), which suggests a wider importance of *yeaG* and *yeaH* and pathways affected by them in bacterial adaptive processes. Although the product of *yeaH* has no sequence or structural homology to any known protein, YeaG is a Hank’s type kinase (11,12). In earlier work, we revealed that the viability of *ΔyeaG Escherichia coli* becomes significantly compromised only after 24 h under N starvation and thus proposed a role for YeaG in the adaptive response to sustained N starvation (6). Further, we showed that the intracellular concentration of RpoS (σ^38^), the major bacterial sigma (o) factor required for the transcription of general stress response genes, was decreased by more than 2-fold in *ΔyeaG* bacteria compared to the wild-type (6) and consequently suggested that *ΔyeaG* bacteria might be compromised to effectively adapt their transcriptional programme to survive a sustained period of time under N starvation. In the present study we explore the molecular basis by which YeaG benefits *E. coli* in efficiently adapting to sustained periods under N starvation.

## RESULTS AND DISCUSSION

### The initial 24 h of N starvation represents a physiologically important period during which YeaG-mediated adaptation to sustained N starvation occurs

To decipher the molecular basis by which YeaG contributes to adaptation to sustained N starvation, we began our study by measuring the intracellular levels of YeaG in whole cell extracts from cells experiencing N starvation over 5 days. Results revealed that YeaG levels peaked within the first 24 h upon entry into N starvation, but subsequently diminish (Fig. 1A). This suggests that the YeaG-mediated adaptive response to N starvation occurs, at least under our conditions, within the first 24 h upon entry into the N starvation, despite YeaG expression being induced immediately upon entry into N starvation. Further, since the viability of *ΔyeaG* bacteria gradually decreased only after 24 h under N starvation (N-24) (resulting in ~13.5% of the population at N-24 surviving after 5 days under N starvation, whereas ~62% of the wild-type population remained viable) (Fig. 1B), the results imply that the initial 24 h of N starvation represents a physiologically important period during which YeaG-mediated adaptation to sustained N starvation happens.

**FIGURE 1.**
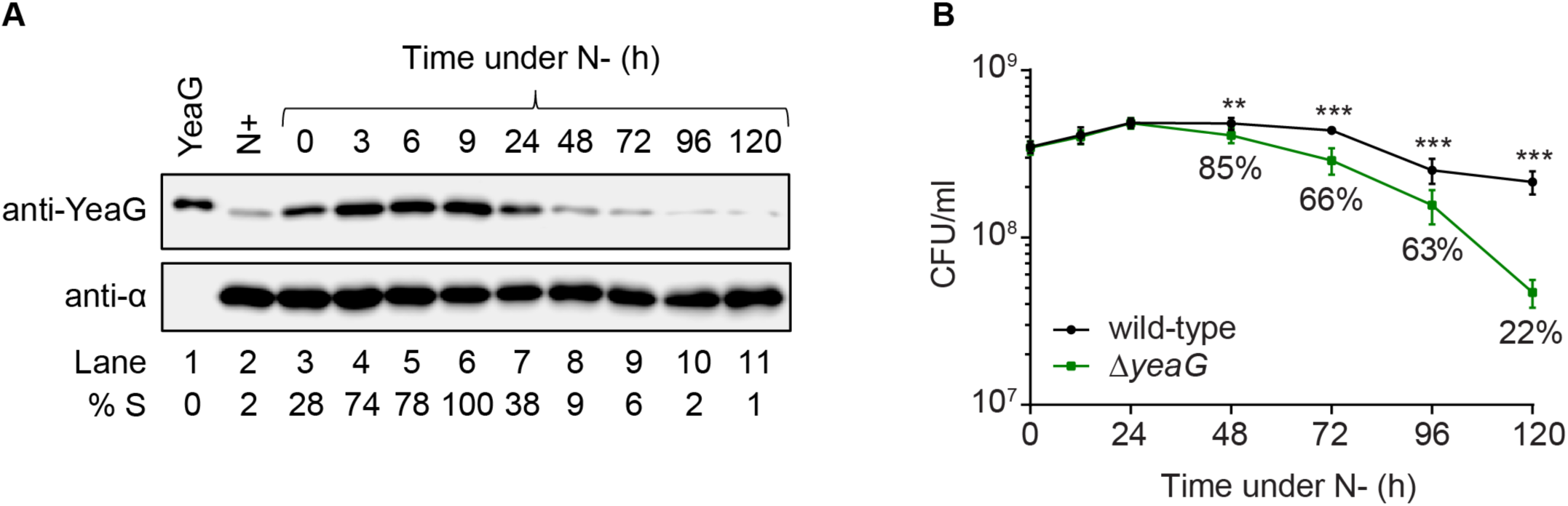
The initial 24 h of N starvation represents a physiologically important period during which YeaG-mediated adaptation to sustained N starvation occurs. (**A**) Representative immunoblot of whole-cell extracts of bacterial cells sampled at N+ and after various periods of N starvation probed with anti-YeaG antibody and anti-RNA polymerase a subunit antibody (loading control). Percentage S (% S) indicates the ratio in intensity between the bands corresponding to YeaG and the a subunit at each time point. (**B**) Graph showing the viability of wild-type and *ΔyeaG* bacteria as measured by counting colony-forming units (CFU) over five days under sustained N starvation. Values show the percentage of viable bacteria in the *ΔyeaG* mutant compared with the wild-type at each time point. Error bars represent s.e.m. (n=3). Statistical analyses were performed by one-way ANOVA (** *p* < 0.01, *\*\*\*p <* 0.001).

### YeaG is required to efficiently shutdown NtrC-activated genes during sustained N starvation

We next compared the transcriptomes of wild-type and *ΔyeaG* bacteria during exponential growth under N replete conditions (N+), following the initial response to N starvation (20 min following N run-out; N-) and after experiencing 24 h of N starvation (N-24) to understand how YeaG contributes to adaptation to sustained N starvation. We defined differentially expressed genes as those with expression levels changed ≥2-fold with a False-Discovery-Rate adjusted *p*-value <0.05. The initial transcriptional response to N starvation involves the activation of genes required for scavenging of alternative N sources by NtrC (4,5). We therefore compared how the transcript levels of NtrC-activated genes (as defined in (5)) differed in wild-type and *ΔyeaG* bacteria at N-(compared to N+) and N-24 (compared to N-). Results shown in Fig. 2A and Fig. 2B reveal that the expression of the NtrC-activated genes at N-(compared to N+) did not significantly differ between wild-type and *ΔyeaG* bacteria (also see Fig. 2C later). Although the transcript levels of NtrC-activated genes decreased at N-24 (compared to N-), we note that, in both wild-type and *ΔyeaG* bacteria, genes associated with the transport of ammonium *(amtB, glnK)*, glutamine *(glnH, glnP* and *glnQ)* and arginine transport *(argT)* and catabolism *(astA, astB, astC, astD* and *astE)* become significantly more downregulated at N-24 compared to other NtrC-activated genes (compare Fig. 2A and 2B). Thus, it seems that *E. coli* experiencing sustained N starvation shutdown cellular systems required for transport of alternative N sources. Comparison of relative transcript levels of NtrC-activated genes in wild-type and *ΔyeaG* bacteria at N+, N- and N-24 indicates that the general downregulation of NtrC-activated genes is less effective in Δ*yeaG* bacteria compared to wild-type bacteria (Fig. 2C). Notably, the transcript levels of *amtB, flgM* (anti-σ^28^ factor; see below) and *yeaH* (see below) were higher in Δ*yeaG* bacteria than in wild-type bacteria at N-24, N- and N+, respectively. Overall, it seems that YeaG has a role in the efficient shutdown of NtrC-activated genes in response to sustained N starvation.

**FIGURE 2.**
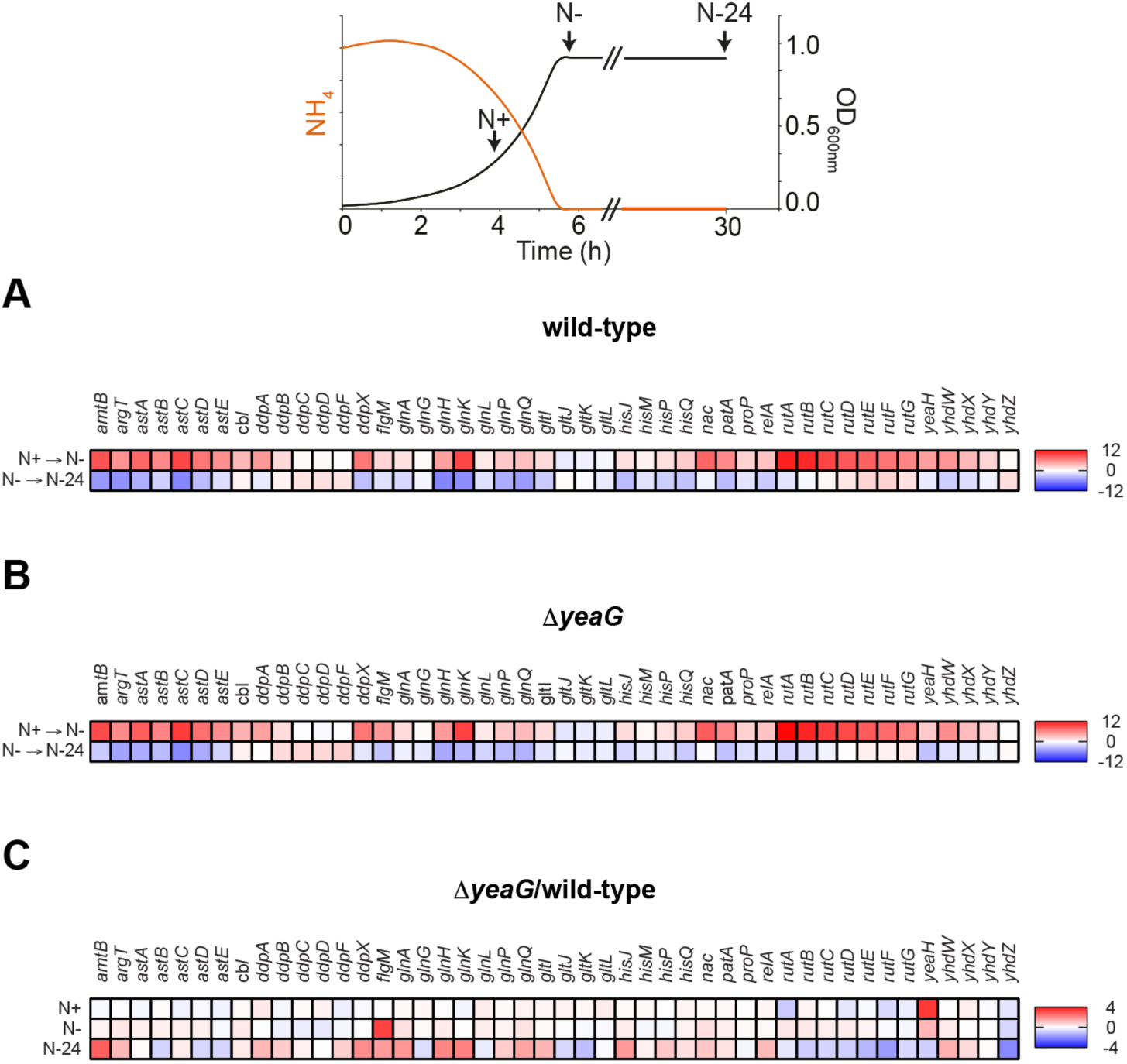
YeaG is required to efficiently shutdown NtrC-activated genes during sustained N starvation. Heat maps showing the expression levels of the 46 genes within the NtrC regulon (as described in (5)) in (**A**) wild-type and (**B**) *ΔyeaG* bacteria at N- (relative to N+) and N-24 (relative to N-) and (**C**) in *ΔyeaG* relative to wild-type bacteria at N+, N-, and N-24. The colour keys on the right indicate the range in fold-change in relative gene expression for each comparison. The schematic representation of the bacterial growth curve at the top indicates the time points: N+, N- and N-24 with respect to the N consumption ([NH4]) and growth (OD_600nm_).

### A role for YeaG in regulating motility-associated gene expression during the initial adaptive response to N starvation

We generated volcano plots to compare the complete transcriptomes of wild-type and Δ*yeaG* bacteria at N+, N- and N-24 (Fig. 3A, 3B and 3C). A total of 21 and 199 genes were differentially expressed in Δ*yeaG* bacteria compared to the wild-type at N- and N-24, respectively (Fig. 3B and 3C, and Table S3 and S4). In contrast, only one gene, *yeaH*, was differentially expressed at N+ in Δ*yeaG* bacteria compared to the wild-type bacteria (Fig. 3A). Among the differentially expressed genes at N- and N-24, we did not detect *yeaH* (Fig. 3B and 3C and Table S3 and S4), suggesting that the properties of Δ*yeaG* bacteria described below are solely due to the absence of *yeaG* and not the dysregulation of *yeaH* expression (also see Fig. S3 later). Intriguingly, although Δ*yeaG* bacteria contain significantly reduced intracellular levels of σ^38^ under sustained N starvation (6), only a small subset (none at N-, and ~8% at N-24) of the σ^38^-regulon (as defined by (8)) was differentially expressed in Δ*yeaG* bacteria, suggesting that the dysregulation of expression of the σ^38^ regulon is unlikely to contribute to the inability of the Δ*yeaG* bacteria to survive sustained N starvation (Fig. S1). Conversely, categorisation of differentially expressed genes using Clustering of Orthologous Groups (COG) analysis (18) revealed that the majority of upregulated genes at N- in Δ*yeaG* bacteria were associated with the biosynthesis of flagella and motility (*fliC, fliD, motA, motB, tap, tar, tsr, fimI*, and *flgK*) (Fig. 3D and Fig. S2). Spinning of *E. coli* cells is indicative of the presence of functional flagella when tethered to a glass surface. Therefore, we examined samples of Δ*yeaG* and wild-type bacteria from N- by phase-contrast microscopy. As shown in Movie S1, we failed to detect any spinning cells in the wild-type sample. In contrast, the Δ*yeaG* sample contained several spinning and motile cells (Movie S2). Interestingly, however, in Movie S2 we note that only some of the Δ*yeaG* cells were spinning or motile, while most cells resembled the wild-type cells, suggesting that the upregulation of motility genes in Δ*yeaG* population at N- could occur in a heterogeneous manner. We further note that the motility-associated genes in Δ*yeaG* bacteria are down-regulated to near-wild-type levels at N-24 (Fig. S2), suggesting that dysregulation of these genes in Δ*yeaG* bacteria is a property of mutant bacteria that is specific to the initial adaptive response to N starvation. We thus conclude that YeaG could have a role in shutting down motility-associated gene expression during the initial adaptive response to N starvation.

**FIGURE 3.**
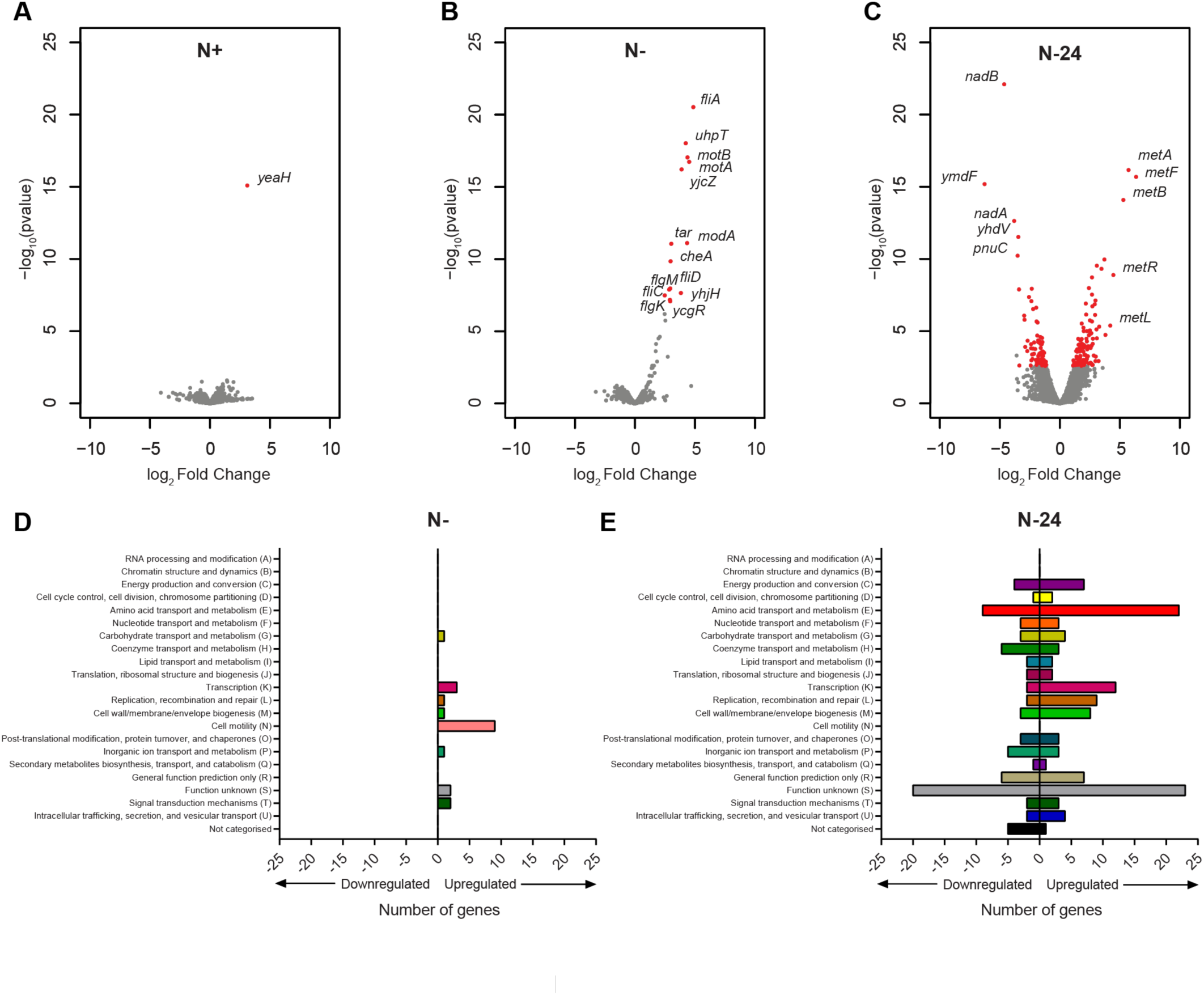
YeaG is required for regulation of the transcriptional response to initial and sustained N starvation. **(A-C)** Volcano plots revealing expression levels of all genes at N+, N- and N-24 (for A, B and C respectively) in *ΔyeaG* bacteria as a fold change from wild-type, where red indicates differentially expressed genes changed ≥2-fold with a False-Discovery-Rate adjusted P-value <0.05. The identities of genes that are most differentially expressed genes and relevant to the study are provided and shown in red. (**D**) Graph categorising all differentially expressed genes at N- based on their COG annotation. (**E**) As in (D) but for N-24.

### A role for YeaG in regulating methionine biosynthesis genes in E. coli experiencing sustained N starvation

Since the viability of the Δ*yeaG* population begins to decline after 24 h under N starvation, we next focused on the differences in the transcriptomes of wild-type and Δ*yeaG* bacteria at N-24. COG analysis of differentially expressed genes at N-24 revealed that the majority of the upregulated genes were either uncharacterised or associated with amino acid transport and metabolism (Fig. 3E), and of the latter group, genes involved in the biosynthesis of methionine were most highly upregulated (Fig. 3C and Table S4). Conversely, genes associated with coenzyme transport and metabolism, and particularly NAD^+^ biosynthesis, were most highly downregulated (Fig. 3C and Table S4). Interestingly, aspartate serves as the precursor for both NAD^+^ and methionine biosynthesis, thus the data suggests that aspartate becomes preferentially directed towards methionine biosynthesis in Δ*yeaG* cells at N-24. Transcript levels of genes of the MetJ and MetR regulons, which include methionine biosynthesis genes, were downregulated in wild-type and Δ*yeaG* bacteria in the initial response to N starvation (at N-, compared to N+; Fig. 4A and 4B). However, at N-24 (compared to N-) these genes (notably, *metL, metA, metB, metC, metF* and *metK)* become upregulated in wild-type bacteria (Fig. 4A). Strikingly, in Δ*yeaG* bacteria, the magnitude by which this upregulation occurred was substantially enhanced compared to wild-type bacteria (Fig. 4B and 4C and Table S4). Quantification of transcript levels of the most dysregulated genes in *ΔyeaG* bacteria *(artJ, metA, metB, metF, metR, mmuP, nadA, nadB, pnuC*, and *yhdV)* by qRT-PCR in wild-type, Δ*yeaG* bacteria, and Δ*yeaG* bacteria in which *yeaG* was complemented from an inducible plasmid, validated the RNA-seq data and confirmed that the differences in transcript levels were due to the absence of *yeaG* (Fig. S3). Overall, we conclude that YeaG has a role in the transcriptional regulation of methionine biosynthesis genes in *E. coli* experiencing sustained N starvation.

**FIGURE 4.**
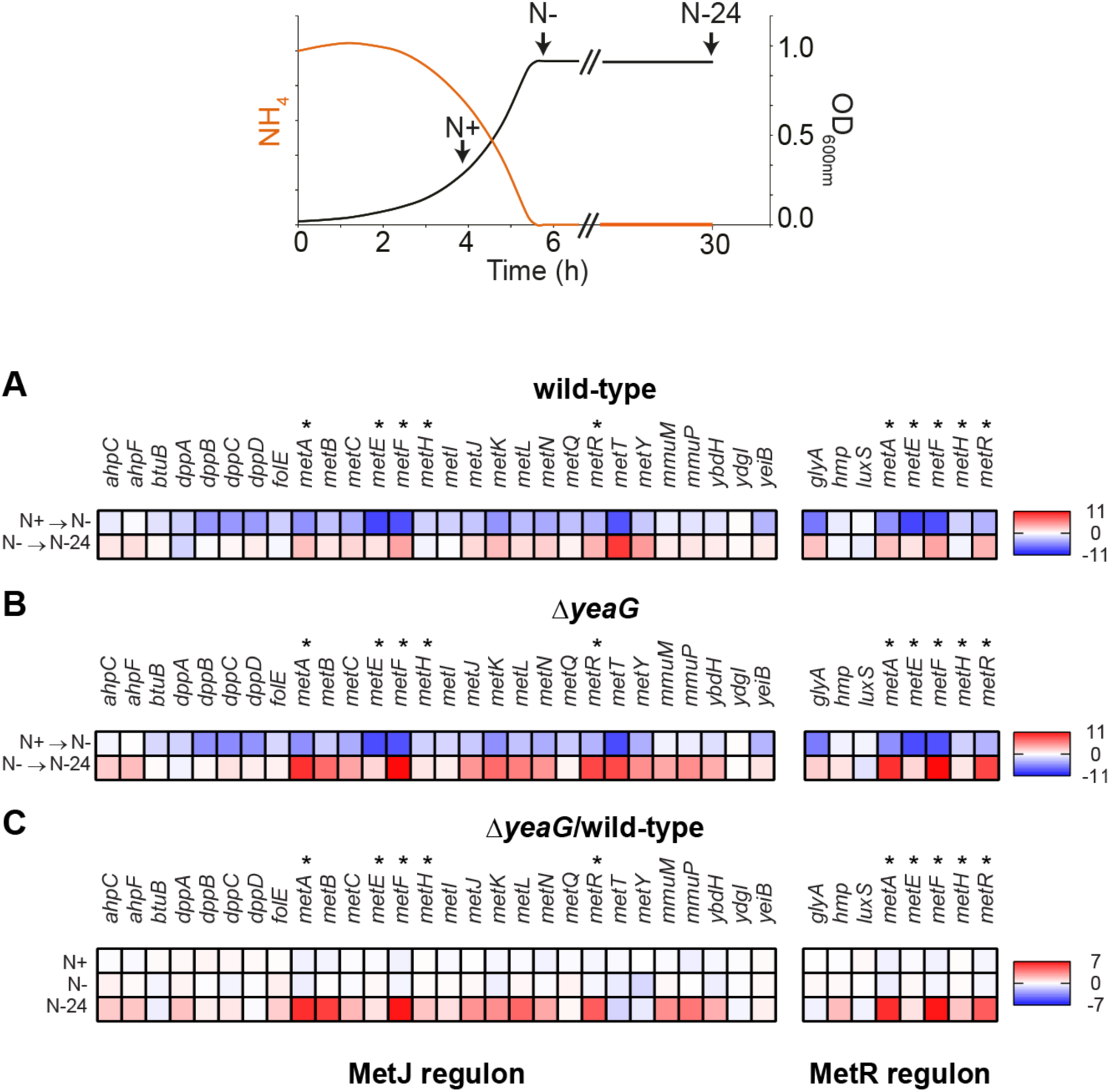
The MetJ regulon is dysregulated in *ΔyeaG* bacteria experiencing sustained N starvation. Heat maps showing expression levels of genes of the MetJ (left) or MetR (right) regulon in wild-type (**A**) and *ΔyeaG* (**B**) bacteria at N- relative to N+, and N-24 relative to N-, and in *ΔyeaG* relative to wild-type bacteria at N+, N-, and N-24 (**C**). The colour keys on the right of each heat map indicate the range in fold-change in relative gene expression for each comparison. Genes which are subjected to regulation by both MetJ and MetR are marked with an asterisk. The schematic representation of the bacterial growth curve at the top indicates the time points: N+, N- and N-24 with respect to the N consumption ([NH_4_]) and growth (OD_600nm_).

### Aberrant activation of the methionine biosynthesis pathway after 24 h in N starved ΔyeaG bacteria

As synthesis of one methionine molecule in *E. coli* requires seven molecules of ATP (19), we considered whether the aberrant synthesis of methionine and associated metabolites of the pathway in Δ*yeaG* bacteria compromises the viability of the mutant population after 24 h under N starvation. We used liquid chromatography-electrospray ionisation mass spectrometry metabolomics to carry out a targeted analysis of selected intermediates (O-succinylhomoserine, methionine, S-adenosylmethionine (SAM) and S-adenosylhomocysteine (SAH)) of the methionine biosynthesis pathway in wild-type and Δ*yeaG* bacteria at N+, N- and N-24. The intracellular levels of the measured metabolites did not significantly differ between the wild-type and Δ*yeaG* bacteria at N+ and N- (Fig. 5). However, at N-, the intracellular levels of the measured nucleotides were overall lower than at N+ (Fig. 5). Consistent with the differences in the transcription of methionine biosynthesis genes between wild-type and Δ*yeaG* bacteria at N-24 (Fig. 3C, Table S4 and Fig. 4), the intracellular levels of O-succinylhomoserine, methionine, S-adenosylmethionine (SAM) and S-adenosylhomocysteine (SAH) were higher in Δ*yeaG* cells than in the wild-type at N-24 (Fig. 5). Overall, the results unambiguously confirm that methionine biosynthesis pathway is active and aberrantly upregulated in Δ*yeaG* bacteria following 24 h of N starvation.

**FIGURE 5.**
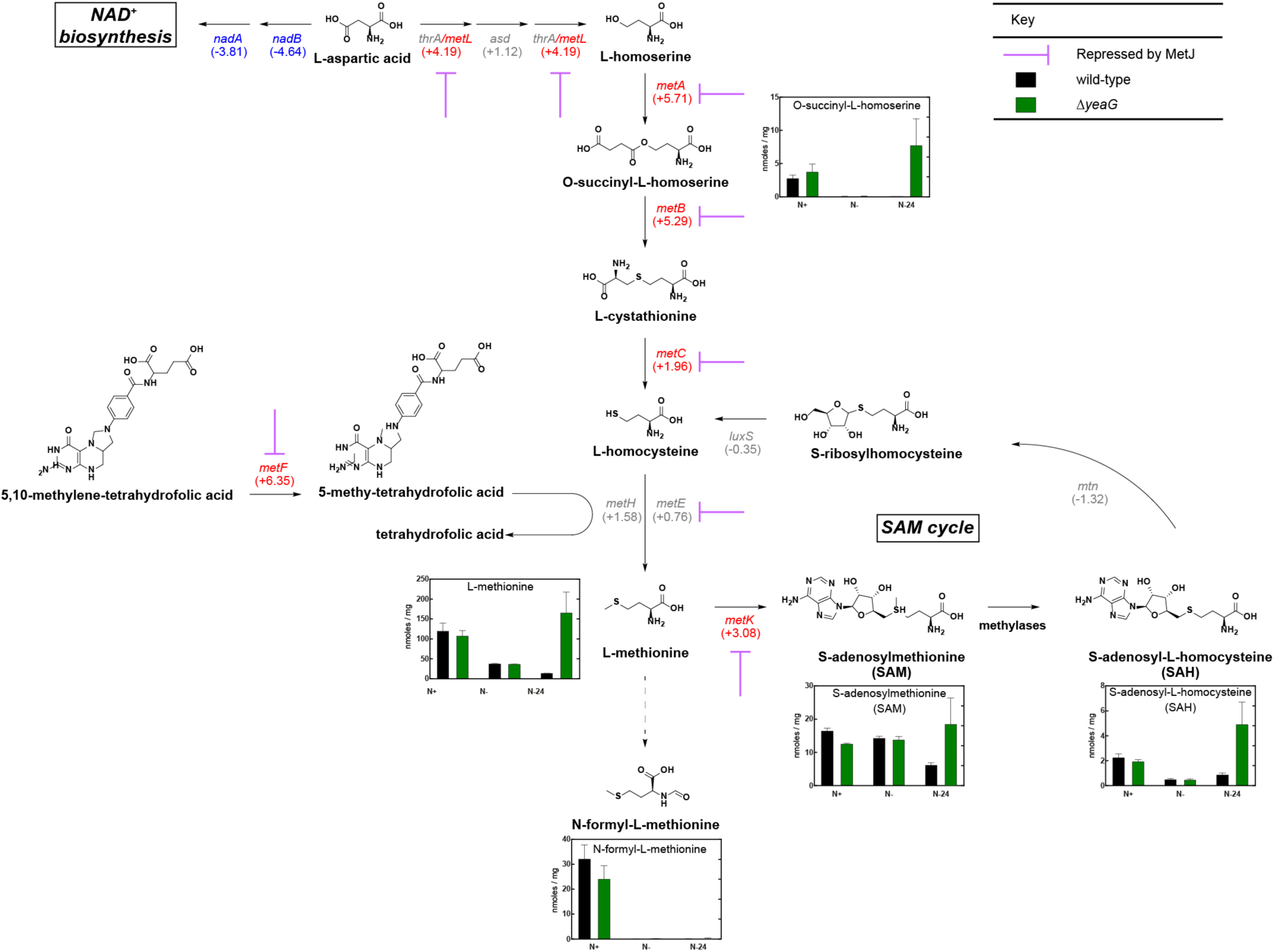
A role for YeaG in regulating methionine biosynthesis genes in *E. coli* experiencing sustained N starvation. Schematic outlining the methionine biosynthetic pathway in *E. coli.* The purple arrows indicate genes that are repressed by MetT. The numbers in parentheses represent the gene expression in *ΔyeaG* relative to wild-type bacteria at N-24 for the associated gene. The concentrations of measured metabolites (nmoles / mg) associated with methionine biosynthesis in either wild-type or *ΔyeaG* bacteria at N+, N- and N-24 are represented in bar charts. Error bars represent s.e.m. (n=3).

### Derepression of transcription leads to activation of the methionine biosynthesis pathway in 24 h N starved ΔyeaG bacteria

Most of the methionine biosynthesis genes that become dysregulated at N-24 in Δ*yeaG* bacteria are negatively regulated by the transcription repressor, MetJ along with its cofactor, SAM, which increases the affinity of MetJ to bind cognate sites by ~100- 1000-fold (20). Therefore, since intracellular SAM levels are ∼3-fold, and MetJ transcript levels are ∼8-fold, higher in Δ*yeaG* cells than in wild-type cells at N-24 (Fig. 5 and Table S4 respectively), it seems paradoxical that MetJ is apparently compromised (leading to upregulation of MetJ-regulated methionine genes) in Δ*yeaG* bacteria. We thus considered whether either the stability or activity (or both) of MetJ might be altered in Δ*yeaG* bacteria under sustained N starvation. To investigate whether the stability of MetJ is affected during the initial 24 h under N starvation in Δ*yeaG* bacteria, we introduced an in-frame fusion encoding three repeats of the FLAG epitope at the 3’ end of MetJ in wild-type and Δ*yeaG* bacteria. To first confirm that the presence of the FLAG epitope did not adversely affect the activity of the MetJ protein, we compared the relative levels of *metF* transcripts (a gene which contains five MetJ binding sites (met boxes) and thus is heavily repressed by MetJ in the presence of excess methionine; Fig. 6A) in wild-type, *ΔmetJ, metJ-FLAG* and Δ*yeaG metJ-FLAG* strains grown in LB broth supplemented with 100 μg/ml methionine. As shown in Fig. 6B, the FLAG epitope did not detectably affect the activity of MetJ. We then used an antibody against the FLAG epitope to measure the relative intracellular levels of MetJ-FLAG protein in whole-cell extracts prepared from wild-type, *ΔmetJ, metJ-FLAG* and Δ*yeaG metl-FLAG* cells as a function of time (up to 24 hours) under N starvation. Results shown in Fig. 6C reveal that the intracellular levels of MetJ-FLAG protein in wild-type, *ΔmetJ, metJ*-FLAG and *ΔyeaG metl*-FLAG cells do not sufficiently differ to warrant the increased levels of *metF* transcription (and by inference, transcription of other MetJ controlled genes) in the Δ*yeaG* bacteria at N-24.

**FIGURE 6.**
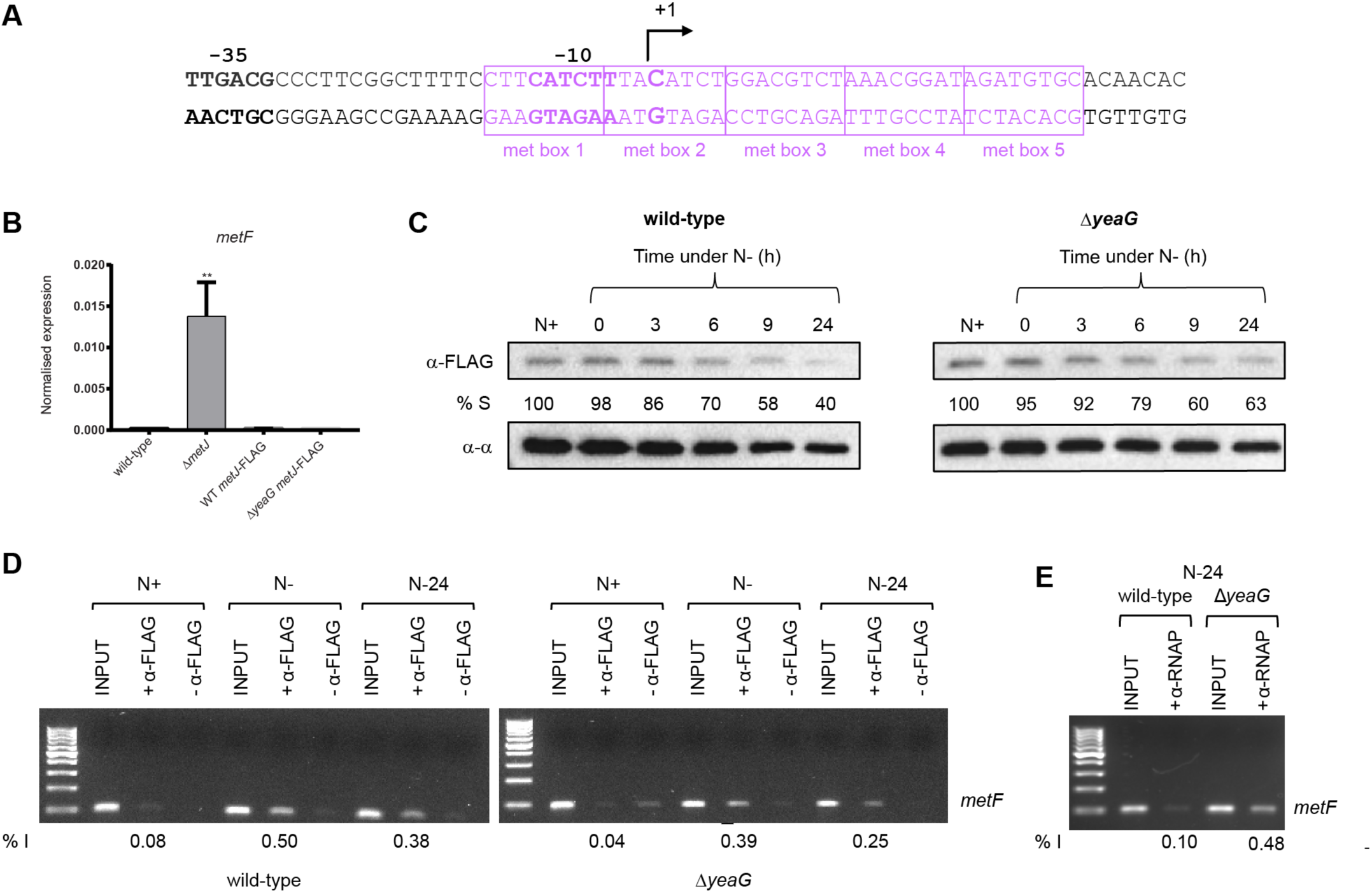
Aberrant activation of the methionine biosynthesis pathway after 24 h in N starved Δ*yeaG* bacteria. (**A**) The sequence of the *metF* promoter showing the consensus -35 and -10 promoter regions with respect to the transcription start site (+1). Boxed in purple are the met boxes. **(B)** Graph showing qRT-PCR results of relative abundance of *metF* transcripts in wild-type, *ΔmetJ, metJ-FLAG* and *ΔyeaG metJ-FLAG* cells normalised to 16S internal control (see text for details). Error bars represent s.e.m. (n=3). Statistical analyses were performed by one-way ANOVA (** *p* < 0.01). **(C)** Representative immunoblot of whole-cell extracts of bacterial cells sampled at N+ and following various periods of N starvation probed with anti-FLAG antibody (to detect MetJ) and anti-RNA polymerase alpha subunit antibody (loading control). Percentage S (% S) indicates the ratio in intensity between the bands corresponding to MetJ and the alpha subunit at each time point. **(D)** Representative image of an agarose gel showing PCR products corresponding to the *metF* promoter region in the N+, N- and N-24 samples immunoprecipitated with anti-FLAG antibodies i.e. bound by MetJ (see text for details). **(E)** As in (D) but anti-β antibodies were used to immunoprecipitate RNAP bound to the *metF* promoter region at N+, N- and N-24. In (D) and (E), % I indicates band intensity of the signal corresponding to the *metF* promoter region relative to the input positive control sample.

We next investigated whether the DNA binding activity of MetJ is adversely affected in Δ*yeaG* bacteria at N-24. To do this, we used MetJ binding to the *metF* regulatory region as a proxy to investigate the DNA binding activity of MetJ *in vivo* by chromatin immunoprecipitation (ChIP). We sampled *metJ*-FLAG and Δ*yeaG metl*-FLAG cells at N+, N- and N-24, immunoprecipitated MetJ-FLAG bound DNA fragments using anti-FLAG antibodies, and used rate-limiting semi-quantitative PCR for specific enrichment of the *metF* regulatory region. As shown in Fig. 6D, the amount of MetJ binding to the *metF* regulatory region was decreased by ~1.3 (at N-) and ~1.5 (at N-24) fold, suggesting that the DNA binding activity of MetJ is slightly compromised in *ΔyeaG* bacteria compared to wild-type bacteria. Since the met boxes at the *metF* promoter overlap the consensus -10 promoter region (Fig. 6A), it is conceivable that repression of transcription by MetJ at the *metF* promoter occurs by occlusion of RNA polymerase (RNAP) binding to the promoter. Therefore, given the compromised ability of MetJ in *ΔyeaG* bacteria to bind to the *metF* promoter region (Fig. 6D), it is conceivable that MetJ is outcompeted by RNAP in *ΔyeaG* bacteria, resulting in the enhanced transcriptional derepression of *metF* and, by inference, other MetJ regulated genes. To investigate this, we immunoprecipitated RNAP and repeated the experiment described in Fig. 6D. As shown in Fig. 6E, an ∼4.8-fold increase in RNAP binding to the *metF* promoter was detected in *ΔyeaG* bacteria compared to wild-type bacteria at N-24. Overall, we conclude that the aberrant upregulation of methionine biosynthesis genes in *ΔyeaG* bacteria at N-24 occurs because of derepression of transcription of these genes.

## CONCLUSIONS

Many bacterial adaptive responses begin with, and are accompanied by, large-scale changes to the transcriptome. The coordination and the scale of the changes that occur in the transcriptome are often underpinned by multiple and complex control layers which ensure the appropriateness of the adaptive response to benefit the cell. The initial adaptive transcriptional response to N starvation in *Escherichia coli* and related bacteria is orchestrated by the global transcription factor, NtrC. This initial response, which is energetically costly, allows the cells to adjust their physiology to actively import and assimilate N from alternative resources. How the transcriptome changes and the regulatory basis underpinning such changes during *sustained* N starvation are not well understood. The results of this study show that the NtrC regulon is shutdown in *E. coli* experiencing sustained N starvation and that YeaG contributes to the effectiveness of this shutdown. This intuitive response to sustained N starvation likely serves to prevent energy wastage in the absence of alternative N sources to sustain growth. Further, we have uncovered that YeaG is part of the regulatory sub-circuit controlling the expression of methionine biosynthesis genes in sustained N starved *E. coli.* The results suggest that the aberrant and energetically costly activation of the methionine biosynthesis pathway has detrimental effects on the viability of the *ΔyeaG* bacteria. It appears YeaG contributes to the efficiency by which MetJ-repressed genes are silenced in *E. coli* experiencing sustained N starvation. Thus, the absence of YeaG leads to the transcriptional derepression of these genes. Although the initial adaptive response to N starvation is largely a scavenging response and does not involve chemotactic-like behaviour, intriguingly, some cells in the *ΔyeaG* population synthesise functional flagella and display motility upon entry into N starvation. This observation implies a potential role for YeaG in regulating flagellum biosynthesis genes and motility during the initial adaptive response to N starvation, which warrants further investigation. Thus, since it seems that potentially two distinct systems (flagellum and methionine biosynthesis) are affected by YeaG in response to N starvation, we speculate that YeaG (recall that it is a kinase) could have more than one phosphorylation target in *E. coli.* Overall, the transcriptional responses of *ΔyeaG* bacteria in the N starved state suggest that YeaG acts as a ‘molecular brake’ in *E. coli* experiencing N starvation. Further, we note that the intracellular levels of SAM are ∼3-fold higher in *ΔyeaG* bacteria compared to wild-type bacteria at N-24. SAM is the second most abundant cofactor (behind ATP), and an important nucleoside serving as a methyl donor in a broad array of metabolic and biosynthetic reactions, including methylation of DNA, RNA, proteins and lipids. Methylation can have direct biochemical relevance in metabolic reactions and is involved in transcription regulation at the epigenetic level. Thus, it is tantalising to speculate whether epigenetic control of the transcriptome, through methylation or phosphorylation, happens during sustained N starvation and the role of YeaG in this process. In summary, although the transcriptional responses to transient growth arrests or during growth transitions due to nutrient starvation (e.g. diauxic growth) has been explored, this study uniquely demonstrates that the transcriptome of growth attenuated bacteria experiencing sustained nutrient starvation can be plastic and that this plasticity is subject to regulation.

## EXPERIMENTAL PROCEDURES

### Bacterial strains, plasmids, growth conditions and viability measurements

Strains and plasmids used in this study were derived from *Escherichia coli* K-12 and are listed in Table S1. The *metJ-*FLAG strains were constructed using the λ Red recombinase method to introduce an in-frame fusion encoding three repeats of the FLAG epitope (3 x (DYKDDDDK)) followed by a kanamycin resistance cassette amplified from the pDOC-F plasmid (13), to the 3’end of *metJ.* The *ΔmetJ* strain was also constructed using the X Red recombinase method to introduce an in-frame fusion encoding a kanamycin resistance cassette amplified from the pDOC-K plasmid (13) in place of the *metJ* gene in the wild-type *E. coli* strain. Bacteria were grown in Gutnick minimal medium according to Figueira *et al.* (6), where the sole source of nitrogen is NH_4_Cl; overnight cultures were grown in medium containing 10 mM NH_4_Cl and N-limiting growth curves were carried out in medium containing 3 mM NH_4_Cl. To measure the biological activity of MetJ-FLAG, the *metJ-*FLAG strains were grown in LB (Luria-Bertani) medium supplemented with 100 μg/ml methionine to induce repression of the methionine biosynthesis pathway as described in (14). Samples were taken for analysis of *metF* transcript levels as described below by quantitative real-time PCR taken during exponential growth. All growth curves were generated by measuring the optical density (OD_600nm_) of the bacterial cultures over time. The number of viable cells was determined by measuring colony forming units (CFU) ml^-1^ from serial dilutions on LB agar plates at the designated time points shown in Fig. 1.

### Antisera and immunoblotting

Immunoblotting was completed according to standard laboratory protocols (15). To produce an antiserum against YeaG, His-tagged YeaG (expressed from the arabinose-inducible plasmid: pBAD18-*yeaG* in MC1061) was purified using nickel affinity purification. Eurogentec produced the antiserum against YeaG through immunisation of rabbits with the purified YeaG protein. This protein was also used to affinity-purify the antibodies from the sera following standard laboratory protocols (15). The YeaG antibody was used at 1:2,500 dilution. The following commercial primary antibodies were used: mouse monoclonal anti-RNAP α subunit 4RA2 at 1:10,000 dilution (Biolegend), and anti-FLAG M2 at 1:1,000 dilution (Sigma, F3165). Both secondary antibodies were used at 1:10,000 dilution, either rabbit anti-mouse IgG H&L horseradish peroxidase (HRP)-linked secondary antibody (Abcam, ab97046) or Amersham ECL Rabbit IgG, HRP-linked whole antibody (from donkey) (GE Healthcare, NA934). ECL Prime western blotting detection reagent was used for all the immunoblotting experiments done in this study (GE Healthcare, RPN2232).

### RNA sequencing (RNA-seq)

Cultures were grown as in (6) and sampled at the following time points: N+, when the OD_600_nm = 0.3; N-, 20 minutes following N runout; and N-24, where bacteria were subjected to 24 hours of N starvation. Two biological replicates were taken for N+ and N-, and three for N-24. Cultures were mixed with a 1:19 phenol:ethanol solution at a ratio of 9:1 culture:solution and then pelleted. Cell pellets were sent to Vertis Biotechnologie AG for downstream processing exactly as previously described by (16). The data analyses were performed with the ‘CLC Genomics Workbench 7 using standard parameters. RNA-seq reads were mapped to the *E. coli* K-12 MG1655 (U00096) genome using Burrows-Wheeler Aligner. Reads that mapped uniquely were used for further analysis. The number of reads mapping to each gene was calculated and matrix of read counts was generated. The matrix was analysed using the DESeq2 BioConductor package for differential gene expression analysis. Genes with ≤10 reads mapped to them were excluded from analysis. All statistical analyses were performed in R version 3.4.2.

### Quantitative real-time PCR (qRT-PCR)

RNA was acquired and stabilised at specified time points using Qiagen RNA Protect reagent (Qiagen, 76526). RNA extraction was carried out using the PureLink RNA mini kit (Invitrogen, 12183025) and DNase Set (Invitrogen, 12185010). The High-Capacity cDNA Reverse Transcription kit (Applied Biosystems, 4368814) was used to convert 100 ng of purified RNA to cDNA. 400 ng/μl of cDNA was used per qRT-PCR reaction containing: 10 μl PowerUp SYBR Green Master Mix (Applied Biosystems, A25742), 1 μl forward primer (5 μM), 1 μl reverse primer (5 μM), and 1 μl template cDNA in a 20 μl reaction volume. Amplifications were performed on the StepOnePlus Real-Time PCR system (Applied Biosystems, 4376600) using the following cycle: 95 °C (10 mins) followed by 40 cycles of 95 °C (15 s), 59 °C (15 s), 68 °C (45 s), and a melt curve of 95 °C (15 s), 60 °C (1 min), 95 °C (15 s). Sequences of all primers used in this study are listed in Table S2. All real-time analysis was performed in triplicate and compared against 16S expression as an internal control.

### Metabolomics

Metabolite analysis was carried out using liquid chromatography mass spectrometry (LC-MS) based metabolomics. Briefly, cultures of *ΔyeaG* and WT strains were grown in 100 ml of Gutnick minimal medium in triplicates as described above for the RNA-seq experiment. The cultures were then spun down rapidly at 3,000 xg for 5 min, the excess of the supernatant was discarded and the remaining 10 ml of the culture were passed through a 0.22 pore size filter (mixed cellulose ester membrane; Millipore GSWP02500). The metabolites were then quenched by submerging the filter containing the bacteria directly into 1 ml solution of acetonitrile: methanol: water (ACN:MeOH:H_2_O; 2:2:1 v/v/v) that was pre-chilled at -40 °C on dry ice. The whole mixture was then transferred into a 2 ml screw cap polypropylene tube containing around 400 μl of acid-washed beads (150-212 μm; Sigma G1145) and ribolysed for 40 sec at 6.5 speed. Following lysis, the tubes were centrifuged at 12,000 rpm for 10 min and the supernatants containing soluble metabolites were passed into a Spin-X 0.22 μm cellulose acetate centrifuge tube filter (Costar, 8160) and stored at -80 °C. Prior to LC-MS, metabolite samples were mixed with acidified ACN (0.2% acetic acid) at 1:1 ratio, spun down at top speed for 5 min and then transferred into LC-MS vials. LC-MS was performed in an Agilent 1200 LC system coupled with an Agilent Accurate Mass 6230 TOF apparatus using the method as described in (17). Metabolite standard samples of O-succinylhomoserine ([M+H]+ m/z = 220.0822, RT = 7.77), methionine ([M+H]+ m/z = 150.5609, RT = 8.28), S-adenosylmethionine (SAM) ([M+H]+ m/z = 400.1497, RT = 17.1), S-adenosylhomocysteine (SAH) ([M+H]+ m/z = 386.3493, RT = 13.36) and N-formyl-L-methionine ([M+H]+ m/z = 178.0543, RT = 1.33) were also run alongside at a range of concentrations. Metabolite abundances and concentration calculations were performed using Agilent’s Mass Hunter suite and Mass Profinder software. Metabolite abundance were normalised with residual protein of each sample, which was estimated using BCA protein assay kit (Pierce, 23225).

### Chromatin immunoprecipitation (ChIP)

Cultures were grown in 3 mM NH_4_Cl Gutnick minimal medium as above; 25 ml was sampled at N+, N-and N-24, and processed exactly as previously described by (5). Either anti-FLAG (M2) or anti-β (WP002) antibodies were used for the MetJ-FLAG ChIP and RNAP ChIP, respectively. Primers used for rate limiting semi-quantitative PCR are listed in Table S2.

### Statistical analysis

Unless otherwise specified, all data show the mean average of at least three independent experiments, where variation shown is the standard error of the mean (s.e.m.). Statistical significance was determined using the Student’s t-test or one-way analysis of variance (ANOVA) where a probability (*p*) value of < 0.05 was considered statistically significant (* *p* < 0.05, ** *p* < 0.01, *** *p* < 0.001, ns – not significant).

## ACKNOWLEDGEMENTS

A Wellcome Trust Investigator award WT100958MA and an MRC doctoral studentship funded this work. Work in the L.P.S.C laboratory is supported by the Francis Crick Institute, which receives its core funding from Cancer Research UK (FC001060), the UK Medical Research Council (FC001060), and the Wellcome Trust (FC001060). We thank members of the S.W. laboratory for constructive comments on the manuscript.

## AUTHOR CONTRIBUTION

A.S., D.R.B., and S.W. designed the experiments; A.S., D. E., D.R.B. and R.F. performed them, A.S. and D.E. analysed them, A.S. and D.E. analysed data presented in the manuscript, A.S. and S.W. wrote the manuscript, A.S., S.W., D.R.B. and L.P.S.C. edited it, and all authors approved of the final content in the manuscript.

